# Control of meiotic double-strand-break formation by ATM: local and global views

**DOI:** 10.1101/236984

**Authors:** Agnieszka Lukaszewicz, Julian Lange, Scott Keeney, Maria Jasin

**Author notes:** To whom correspondence should be addressed: Scott Keeney Maria Jasin.

## Abstract

DNA double-strand breaks (DSBs) generated by the SPO11 protein initiate meiotic recombination, an essential process for successful chromosome segregation during gametogenesis. The activity of SPO11 is controlled by multiple factors and regulatory mechanisms, such that the number of DSBs is limited and DSBs form at distinct positions in the genome and at the right time. Loss of this control can affect genome integrity or cause meiotic arrest by mechanisms that are not fully understood. Here we focus on the DSB-responsive kinase ATM and its functions in regulating meiotic DSB numbers and distribution. We review the recently discovered roles of ATM in this context, discuss their evolutionary conservation, and examine future research perspectives.

## Introduction

During meiosis, homologous chromosomes pair and undergo reciprocal exchange of DNA via homologous recombination, which is essential for their faithful segregation at the first meiotic division. In addition, meiotic recombination increases genetic diversity within species.^1–3^ Meiotic recombination is initiated during early prophase I with the formation of programmed DNA double-strand breaks (DSBs) by the evolutionarily conserved SPO11 transesterase in complex with TOPVIBL (**Fig. 1A**).^4,5^ SPO11 generates DSBs at many genomic positions to promote homologous chromosome pairing and recombination.^6^

**Figure 1.**
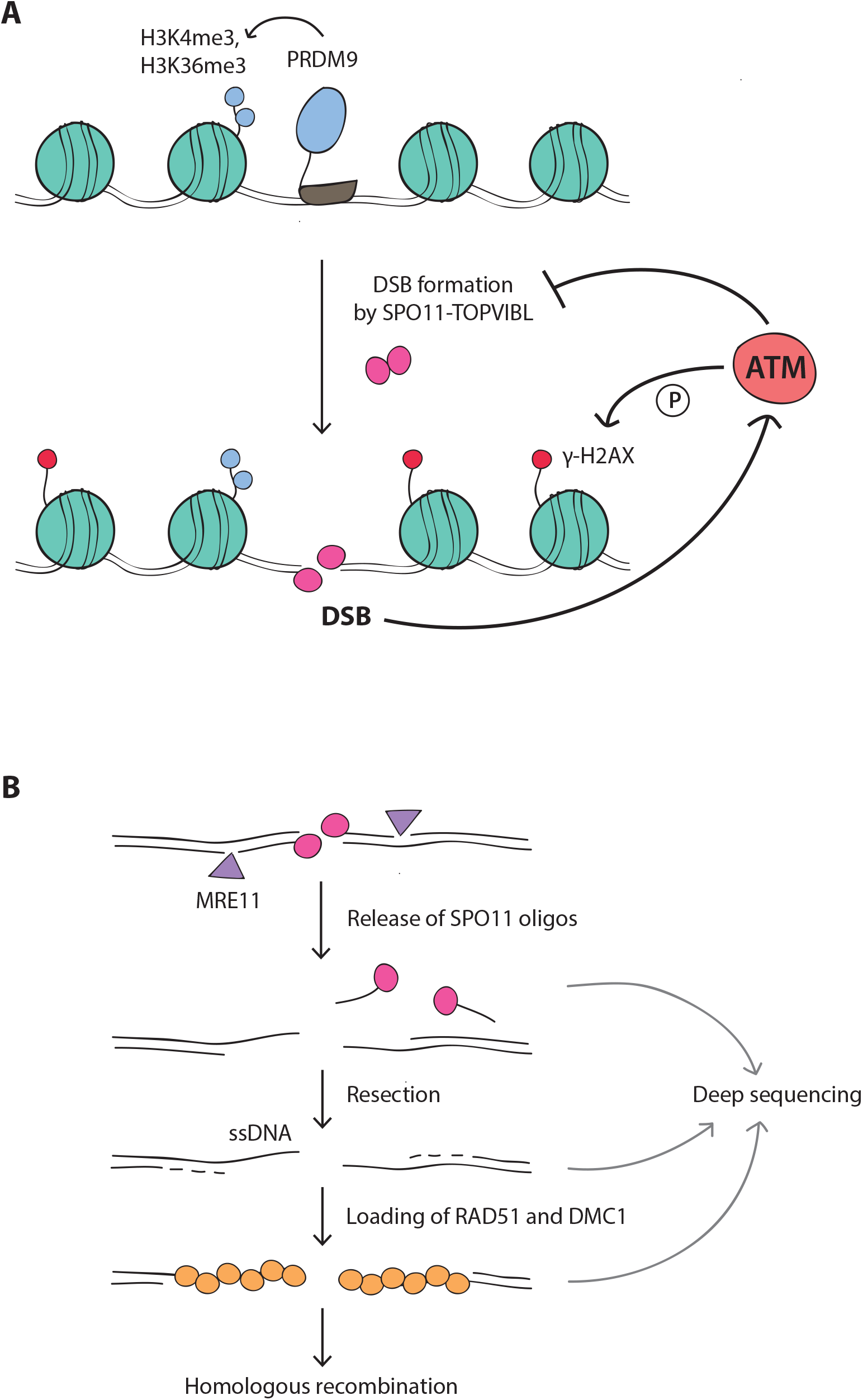
**(A)** Meiotic DSB formation. The SPO11-TOPVIBL complex induces DSBs at genomic locations enriched for H3K4me3 and H3K36me3, histone modifications deposited by PRDM9. DSBs activate the ATM kinase, which suppresses SPO11 from further DSB formation by an as yet unknown mechanism. In response to DSBs, ATM phosphorylates histone H2AX (γ-H2AX) in surrounding chromatin. **(B)** Meiotic DSB processing and detection. SPO11, covalently attached to DNA, is removed by endonucleolytic cleavage by the MRE11 complex. This reaction releases SPO11 bound to short oligonucleotides (SPO11-oligo complexes) and exposes short single-stranded DNA (ssDNA) tails at DSB sites. These tails are further extended by resection and become substrates for DMC1 and RAD51 strand exchange proteins, which mediate homologous recombination. SPO11 oligos, ssDNA, and DMC1/RAD51-coated ssDNA can be analyzed by deep sequencing to map genome-wide DSB positions (see main text for references).

DSBs are dangerous, as failure to repair them can compromise genome integrity.^7–9^ Therefore, meiotic cells control DSB numbers and mostly restrict DSB formation to a narrow window of time during prophase I.^10,11^ Moreover, in most species, meiotic DSBs are not randomly distributed throughout the genome but are primarily (but not exclusively) localized to narrow genomic regions referred to as “hotspots”.^6,12–15^ DSB hotspots themselves are unevenly spread, so that different chromosomal regions manifest as DSB-“rich” and DSB-”poor”.^13,16^ The shape of this “DSB landscape” is molded by a complex web of factors operating over different size scales (from single base pairs to whole chromosomes; expanded further in this review).^16–18^ In addition to its effects on genome diversity and evolution, dictating the position of recombination events likely promotes correct DSB repair.^16,19^ Thus, understanding how meiotic DSBs are numerically and spatially patterned is an exciting and important area of current research.

In recent years, it has become evident that the DNA damage-responsive kinase ATM (<u>a</u>taxia telangiectasia <u>m</u>utated) plays a pivotal role in regulating meiotic DSB formation in various species.^17,20–26^ In mouse, where the lack of ATM causes a particularly extreme excess of DSBs, this deregulated DSB formation appears to be a major cause of meiotic defects in ATM-deficient spermatocytes.^22,27–30^ It has been proposed that ATM, activated by DSBs, operates via a negative feedback loop to restrain SPO11 from further DSB formation.^10,20,22,26,31^ Because ATM is activated in the vicinity of the break, this model predicts that the additional DSBs formed in the absence of ATM will display an altered distribution.^22^ Indeed it was found that the budding yeast ATM ortholog, Tel1, inhibits formation of DSBs nearby one another on the same chromatid^20^ and also prevents cleavage of the same location on the sister chromatid and homolog chromatids.^25^ The Tel1-dependent process of distance-dependent DSB suppression, termed “DSB interference”, has the power to mold the DSB distribution.^12,23^ Thus, ATM/Tell-mediated negative feedback is thought to provide a conserved regulatory mechanism controlling the number and positions of DSBs.

Recent studies of genome-wide DSB distributions in mouse^17^ and yeast^23^ have revealed that the loss of ATM/Tel1 leaves distinct footprints on DSB distributions at different size scales. Here, we focus on research performed in mouse, which has uncovered the prominent role of ATM in shaping the genome-wide DSB distribution.^17^ We discuss this discovery in relation to parallel work in the budding yeast *Saccharomyces cerevisiae* demonstrating that Tel1 contributes to the global DSB distribution in that organism as well.^23^ We also highlight species-specific differences in how the DSB landscape responds to the loss of ATM/Tel1 at shorter and longer size scales. Together, these studies expand our view of the local and global roles of ATM/Tel1 in molding DSB distributions. We present an overview of advances made in understanding the spectrum of ATM/Tell-mediated DSB control and we discuss potential underlying mechanisms.

## ATM-dependent regulation of DSB numbers

The average number of meiotic DSBs per cell varies over a wide range in different species and is not correlated-with genome size or chromosome number, suggesting the existence of species-specific (and thus genetically influenced) “set points”. For example, approximately ~200-250 DSBs are formed on the 20 mouse chromosome pairs and ~140-170 on the 16 budding yeast chromosome pairs, despite a ~225-fold difference in genome size between the two organisms.^6,16,32–34^ On the other hand, meiotic cells of *Arabidopsis* and *Tetrahymena*, similarly experience ~200 DSBs but each has only 5 chromosome pairs.^35,36^ However, DSB numbers also show substantial cell-to-cell variation even within a single individual, suggesting that the number of DSBs has a strong stochastic component.^6,10,32,37,38^ Nevertheless, meiotic cells generally form “just enough” DSBs to mediate homology search between chromosomes while seeming to avoid making “too many,” likely to minimize the risk of failure of DSB repair. This self-organization of meiotic DSB formation rests in large part on a network of negative feedback circuits that regulate SPOll activity.^10^ ATM is key to this regulation.

ATM is a Ser/Thr kinase mutated in the cancer-prone disease ataxia telangiectasia^39^ and is primarily known for its function in the DNA damage response in somatic cells. ATM is activated by the MREll complex, which senses DSBs (for extensive review see ref.^40^). Once activated, ATM phosphorylates a large number of substrates (e.g., H2AX, CHK2) to regulate two interdependent processes: the cellular DNA damage response and DSB repair.^41–44^ In contrast to mitotic cells, ATM is essential to mammalian meiotic cells, as ATM deficiency in humans and mice causes fully penetrant cell death during meiosis, resulting in infertility.^45–47^ Although ATM may exert similar DNA damage signaling and repair functions in meiosis,^27,30,48,49^ it acquires an additional critical role (or co-opts an existing role) in controlling the number of SPOll-induced DSBs.

Testes from *Atm^-/-^* mice display >10-fold higher levels of SPOll-oligonucleotide (oligo) complexes, a quantitative by-product of DSB formation (**Fig. 1B**), than wild-type testes, and this increase is apparent soon as DSBs first appear in *Atm^-/^* juvenile testes.^22^ Notably, *Spoil* heterozygosity in otherwise wild-type mice has little effect on DSB numbers (~20% reduction^32,50^), whereas *Spoil* heterozygosity in an *Atm^-/-^* mutant background (i.e., in *Spo11*^+/−^*Atm^-/-^* mice) reduces levels of SPO11-oligo complexes by half.^22^ Moreover, increasing *Spo11* gene dosage increases SPO11-oligo complex levels in ATM-deficient, but not ATM-proficient, mice.^22^ These findings indicate that ATM keeps DSB numbers within a normal range even when *Spo11* gene dosage is altered. The 2-fold reduction in DSB numbers in *Spo11*^+/-^*Atm^-/-^* mice has a dramatic effect on meiotic progression, such that spermatocytes complete recombination to progress to the end of prophase I.^22,30,51^ Thus, in wild-type meiotic cells, ATM is likely activated by DSBs to trigger a negative feedback loop that inhibits toxic numbers of DSBs (also see ref.^10^).

This mechanism appears to be evolutionarily conserved. In *Drosophila melanogaster* female meiosis, temperature-sensitive mutants of *tefu*, the fly *Atm* homolog, show 1.5-to 3fold elevated γ-H2AV (phosphorylated H2AV) signals in germaria at the restrictive temperature.^21^ H2AV is the equivalent of mammalian H2AX, and is phosphorylated by either ATM or the ATM-related kinase ATR (MEI-41 in flies) in response to meiotic DSBs.^21,52^ Thus, this phenotype in flies is consistent with the role of ATM in the mouse germline.

In *S. cerevisiae, tel1* mutant cells demonstrate a >2-fold increase in Spo11-oligo complexes,^23^ which is consistent with increases seen in recombination locally and genome-wide.^24,25^ Three earlier studies using DNA electrophoresis-based assays in DSB repair-defective mutant backgrounds, i.e., *rad50S* and *sae2*, had also reported higher DSB frequency in *tel1* mutants at several DSB hotspots and on at least one whole chromosome (not all assayed chromosomes showed this tendency).^20,25,53^ However, two other studies using the electrophoresis-based assays reported reduced DSB frequency.^54,55^ These conflicting results likely arise from the need to perform these assays in DSB-repair defective mutants, which may themselves have altered DSB formation.^10,31,34,56^ In addition to their utility in otherwise wild-type cells, Spo11-oligo assays are likely more reliable because complexes display a long lifespan during prophase I and, as by-products of DSB formation, are relatively insensitive to DSB repair kinetics following their formation.^22,56^ Thus, immunoprecipitation of Spo11-oligo complexes allows for a more robust assessment of Spo11 activity than detection of short-lived DSB intermediates in the chromosome. (The DSB lifespan issue, as well as the likely impact of Tel1 itself on that process, is discussed in more detail in ref.^23^).

Collectively, these data support the conclusion that ATM orthologs play a conserved role in regulating DSB numbers in mouse, flies, and budding yeast. Nonetheless, the consequence of ATM/Tel1 loss is different in the three species. Whereas ATM-deficient mouse spermatocytes arrest early in prophase I, *tel1* budding yeast cells produce viable spores. Given the much smaller fold increase in DSB numbers in yeast than in mouse, this quantitative difference is probably sufficient to explain the more severe phenotype in mouse. The observation that lowering DSB numbers by reducing *Spo11* gene dosage leads to partial rescue of meiotic progression in Atm-mutant spermatocytes also supports this interpretation.

However, in *Drosophila* females, *tefu* mutant eggs show hallmarks of DSB-repair defective female-sterile mutants, even though DSBs appear to increase similarly to yeast, and developing mutant embryos arrest. Of note, however, fly ATM is essential for normal cell proliferation as well,^21,57^ so perhaps yeast and flies are differentially responsive to additional DSBs in causing cell cycle arrest. For instance, *Drosophila* female meiosis might possess a lower intrinsic DSB tolerance threshold. Notably, in contrast to mouse and yeast, flies do not rely on DSB formation for homologous chromosome pairing.^58^ Consequently, the number of meiotic DSBs is low (~14 per oocyte), closer to the required number of crossovers.^59^ Alternatively, the fold change in DSB numbers in the absence of fly ATM might be underestimated because of the indirect quantification method used. For example, additional breaks formed nearby on the same molecule might be undetectable by measurements of γ-H2AV. Such multiple cuts might be as toxic (difficult to repair) as a large excess of well-dispersed DSBs. Whether *Drosophila* ATM regulates DSB distributions as in mouse and yeast remains an open question.

## Regulation of DSB distribution by ATM

Control of the DSB distribution may be as important as regulation of DSB numbers in ensuring faithful genome transmission. Simultaneous DSBs at the same location on more than one chromatid could impede homologous recombination by damaging DNA templates needed for repair, while multiple DSBs nearby on the same chromatid carry the risk of deletion of intervening sequences. Further, in organisms like budding yeast and mouse that rely on DSB formation for chromosome pairing,^60^ clustered DSB formation may interfere with the orderly and/or timely pairing of homologous chromosomes.

The final distribution of meiotic DSBs within a cell results from the interplay between factors acting at multiple levels and in multiple combinations.^6,13,14,16–18,61–63^ For example, the binding and/or activity of the DSB machinery are determined by local factors that predispose chromatin to DSB formation (e.g., histone modifications, nucleosome occupancy) in combination with factors that define higher-order chromosome structures (e.g., loop-axis organization, DSB-promoting chromosome structure proteins).^4,6^ Together, these properties of the DSB machinery and its chromosomal substrate have been described as “intrinsic” factors that pattern the DSB landscape. This is in contrast to another group of factors that are thought to act “extrinsically” by forming regulatory circuits to modulate DSB distribution and thus maintain DSB homeostasis.^4,1^°,^12,16^ The ATM/Tell-dependent negative feedback loop is one such extrinsic DSB determinant.^12,17,23^ Below we discuss evidence from mouse and yeast work that ATM/Tel1 contributes to the spatial patterning of DSBs over different size scales.

### Local DSB patterning (fine-scale distribution)

Despite the potential for SPO11 to act essentially anywhere in the genome, DSBs are preferentially formed at hotspots, discrete genomic positions of high DSB-forming potential. Nucleotide-resolution DSB maps obtained by deep sequencing of SPO11 oligos allows the most precise description of DSB hotspot structure,^16,17,23,64^ though, DSB hotspots have been mapped genome-wide by different approaches, including those that measure resected DNA (**Fig. 1B**).^16,34,62,65–71^

In mouse (and humans), most hotspot locations are determined by PRDM9 (**Fig. 1A**), a meiosis-specific histone methyltransferase with DNA binding specificity that marks nucleosomes flanking future DSB sites by tri-methylation of histone H3 on lysine 4 (H3K4me3) and on lysine 36 (H3K36me3).^17,62,63,66,67,69,70,72–81^ Levels of PRDM9-dependent H3K4me3 and H3K36me3 are a moderate predictor of DSB hotspot activity, with variation of histone modification levels accounting for up to ~40% of variation in hotspot activity.^17,62,63,79^ Interestingly, although H3K4me3 also marks active transcription promoters in mice,^82,83^ PRDM9-dependent H3K4me3 generally occurs outside of promoters, in both genic and intergenic regions, and SPO11 rarely cuts H3K4me3-enriched promoters.^67,69^ But in the absence of PRDM9, e.g., in mice with targeted disruption of *Prdm9*^69^ or in dogs, which naturally lack a functional PRDM9,^84–88^ SPO11 preferentially targets H3K4me3-enriched regions that include promoters. This is likely true as well in finches, which like dogs, also lack a PRDM9.^89^

Yeast hotspots are mostly in promoters,^16,90^ and hence are also enriched for H3K4me3.^18,91^ Moreover, a physical link between H3K4me3 and the DSB machinery has been established in budding yeast,^92,93^ and a potentially parallel mechanism may exist in mice.^94,95^ However, H3K4me3 levels in wild-type yeast cells predict hotspot DSB frequencies poorly if at all,^18^ and mutants lacking certain transcription factors experience changes in H3K4me3 levels at specific hotspots that are completely uncorrelated with changes in DSB frequency.^96^

Recent studies have explored the role of ATM/Tel1 in shaping the local distribution of DSBs in and near hotspots. Mouse DSB hotspots, as defined by clusters of SPO11-oligo sequence reads (**Fig. 2A**), have a characteristic width and on average tend to display a prominent primary peak at the hotspot center and weaker secondary peaks at flanking positions.^17^ ATM deficiency affects this hotspot substructure, with an average SPO11-oligo profile that is detectably wider (143 bp in wild type versus 185 bp in *Atm^-/-^*) and has more prominent secondary peaks. At individual hotspots, this pattern can be seen as a wider distribution of SPO11-oligo signals around the hotspot center (**Fig. 2A**). One explanation for this observation is that the activity of SPO11 becomes less biased toward hotspot centers. For example, ATM loss might affect chromosomal features (e.g., presence or positioning of methylated nucleosomes surrounding positions where DSBs form, see **Fig. 1A**) to cause less centered SPO11 activity. Alternatively, ATM might inhibit SPO11 from cutting the same chromatid more than once, as Tel1 does in budding yeast (see below). In this second scenario, multiple DSBs form on the same DNA molecule and spread outward from the normally centrally constrained positions. Both scenarios would imply that the ATM-mediated inhibition of DSB formation operates locally at small size scales encompassing a single hotspot.

**Figure 2.**
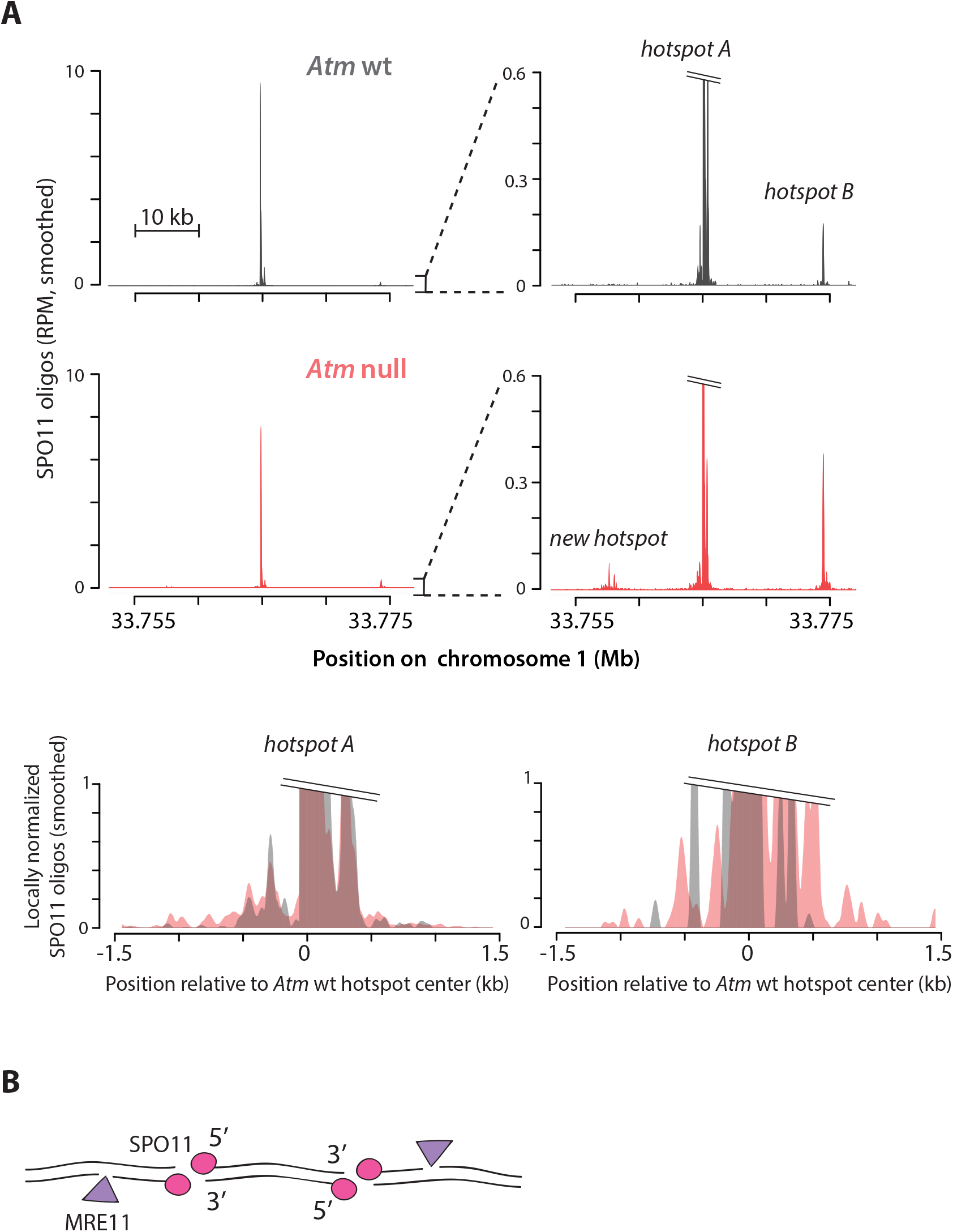
**(A)** Local DSB patterning. (top) Examples of SPO11-DSB hotspots on chromosome 1 in *Atm* wild type (wt) and *Atm* null (data from ref.^17^). Note the emergence of a new hotspot in the absence of ATM. (bottom) Overlays of *Atm* wt and *Atm* null hotspot profiles show a wider distribution of SPO11 oligos around the center of hotspot B in *Atm* null. Methods: As previously described,^17^ hotspots were defined as regions in smoothed SPO11-oligo maps with uniquely mapped reads exceeding 50 times the genome average. The hotspot center was defined as the coordinate of the maximum SPO11-oligo value within the hotspot. Here, SPO11-oligo maps were smoothed with a 101-bp Hann filter. RPM denotes reads per million. **(B)** Schematic of MRE11-independent formation of SPO11 oligos by introduction of nearby DSBs (see main text).

These two possibilities cannot be discriminated by population averaged SPO11-oligo analysis. However, the idea of multiple local cleavages might be partially supported by analysis of SPO11-oligo lengths. Normally, two prominent SPO11-oligo size classes are seen, ~15-27 and ~31-35 nt, but in the absence of ATM this bimodal length distribution is shifted toward longer oligos, leading to a substantial increase in the normally less abundant population of 40-70 nt oligos and an appearance of very long oligos (> 300 nt).^17,22^ These long SPO11 oligos could arise from multiple DSBs formed on the same chromatid, with the 3’ end of a SPO11 oligo created by another DSB nearby rather than MRE11 nicking (ref.^17,23^ and M. Neale, personal communication) (**Fig. 2B**). Thus, the observed alterations in mouse SPO11-oligo lengths and in the SPO11-hotspot width might both reflect multiple DSBs on the same DNA molecule.

Alternatively, but not mutually exclusively, long SPO11 oligos in the absence of ATM could arise from altered end processing (e.g., changes in the positions of endonucleolytic cleavage and/or decreased 3′→5′ exonucleolytic activity by MRE11). In yeast, the length distribution of Spo11 oligos in *tel1* is altered in a fashion similar to those in *Atm^-/^* spermatocytes,^23^ and Tell is know to affect end processing. Tel1 phosphorylates Sae2 and Mrell,^97,98^ which are both required for Spoll-oligo release,^99–103^ and is also important for full-length resection of DSB ends,^104,105^ However, yeast differs from mouse in that individual Spoll-oligo hotspots in *tel1* yeast display a similar width to those in wild type.

In yeast, clear evidence has been provided for Tel1 suppression of multiple DSBs at closely-spaced hotspots on the same chromatid (cis interference).^20^ In the presence of Tel1, the frequency of concurrent DSB formation at hotspots ~≤7.5 kb apart is similar to that expected by chance (no interference), whereas in the absence of Tel1, is higher than expected (negative interference). For example, the absence of Tel1 promotes the appearance of a ~2.4 kb DNA fragment consistent with Spoll cutting at two DSB hotspots within the *HIS4::LEU2* locus. The strength of this Tell-mediated suppression of DSBs decays with distance but can be detected over lengths of 70-100 kb. These observations are consistent with the hypothesis that Tell-mediated suppression primarily functions within a single chromatin loop (~10-20 kb in budding yeast) and to a lesser extent between adjacent loops (discussed in ref.^12^).

Although population averaged Spoll-oligo maps cannot predict DSBs in individual cells, maps in budding yeast provide support for the occurrence of *cis* interference of DSBs: for the two strongest hotspots, *GAT1* and *CCT6*, the neighboring weaker hotspots become stronger when Tel1 is missing.^23^ This phenomenon may also be observed in mouse (**Fig. 2A**), although further analysis of mouse SPOll-oligo maps will be required to reveal similar behavior.

Importantly, Tel1 (and its paralog Mec1) in *S. cerevisiae* also suppresses clustered DSB formation in *trans*, which is the formation of DSBs at the same position on the sister chromatid or homologous chromosome.^20,25^ Because multiple cutting may remove available repair templates or lead to formation of potentially harmful double recombination products, this form of suppression may be important to prevent aberrant meiotic recombination. It has been speculated that the establishment of trans-interference is also locally regulated and distance-dependent, both between sister chromatids and between non-sister chromatids undergoing homologous pairing, although these hypotheses are largely unexplored (discussed in ref.^25,31^).

### Large-scale DSB patterning

At larger size scales, the nonrandom distribution of DSBs defines large chromosomal domains in yeast, mice, and humans.^13,16,17,23,56,64,66,67,90,106,107^ Strikingly, the loss of ATM in spermatocytes leads to substantial large-scale alterations in DSB distribution.^17^ Specifically, large, Mb-scale domains that are normally DSB-poor (cold) appear to be more sensitive to ATM loss than those that are DSB-rich (hot). Consequently, although all SPOll-oligo hotspots become stronger, weaker hotspots that populate DSB-poor domains experience a greater increase in DSB formation. Large-scale ATM-dependent DSB suppression occurs in domains of correlated behavior (measured as the correlation in fold change in SPOll-oligo densities when comparing neighboring regions) and is estimated to decay by ~15-20 Mb. In addition, in the absence of ATM a substantial number of normally weak SPOll-oligo clusters rises above the arbitrary threshold used to call hotspots, which leads to an overall increase in the number of detectable hotspots^17^ (see, e.g., **Fig. 2A**). This behavior predicts that more new hotspots will be detected within normally DSB-poor domains than in DSB-rich domains (**Fig. 3**). Recent findings demonstrate that ATM and PRDM9 contribute independently to hotspot strength.^63^ However, it remains to be elucidated why DSB-poor domains respond more strongly to the absence of ATM and what molecular events within these domains are regulated by ATM to impose this stronger suppression of DSB formation.

**Figure 3.**
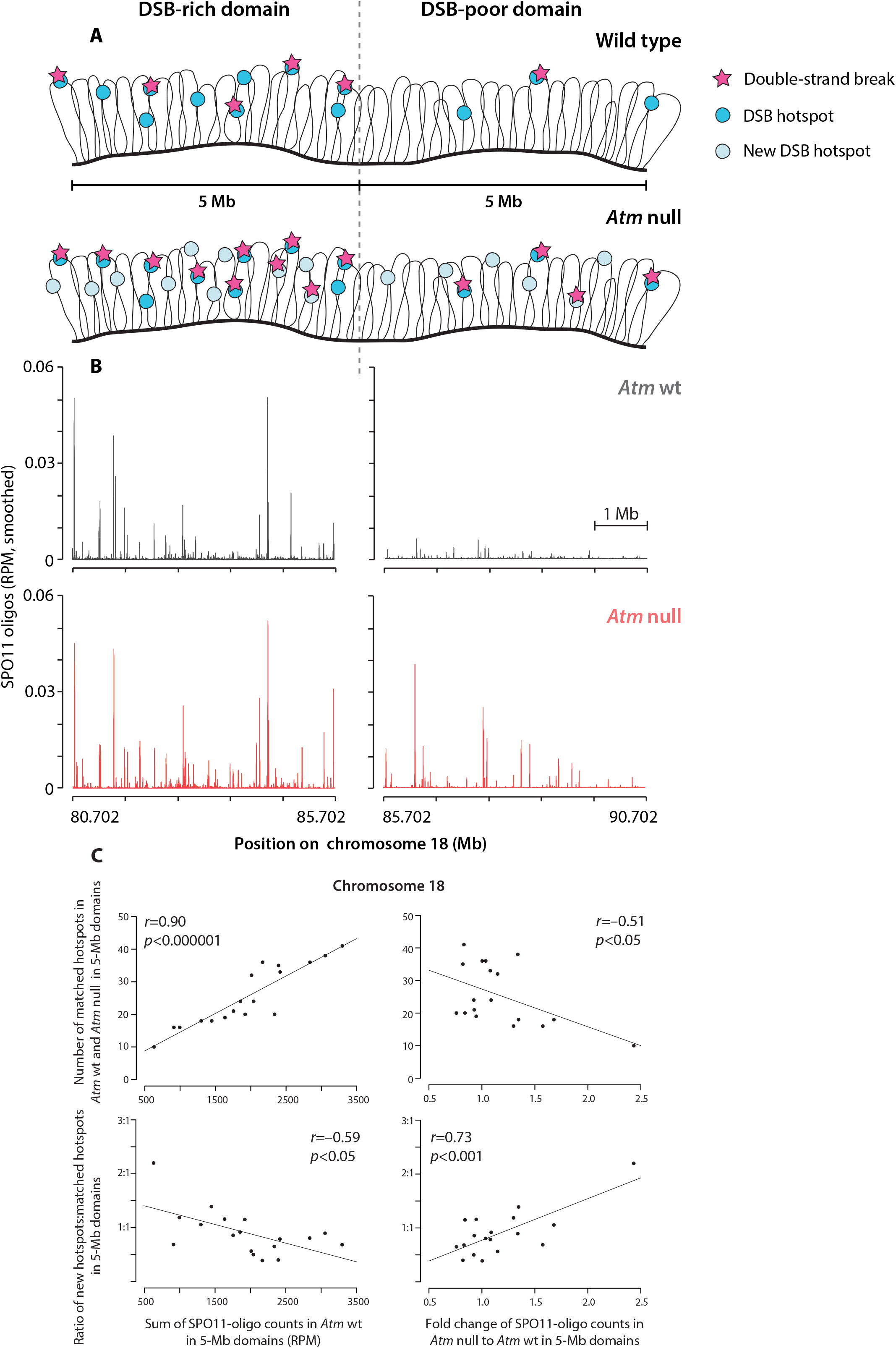
Large-scale DSB patterning. **(A)** Schematic illustrating ATM-mediated DSB distributions across chromosomal domains. (left) Weaker DSB suppression in a DSB-rich domain. In a wild-type cell, this domain contains 10 potential sites for DSB formation (DSB hotspots), of which 5 experience DSBs. In an Atm-null cell, new hotspots emerge (see main text and panel C below), so this domain contains 20 DSB hotspots. Of these, 10 experience DSBs, which gives a 2-fold increase in DSB numbers when ATM is missing. (right) Stronger DSB suppression in a DSB-poor domain. In a wild-type cell, this domain contains 3 DSB hotspots, of which 1 experiences a DSB. In an Atm-null cell, this domain contains 9 DSB hotspots, of which 4 experience DSBs. This gives a 4-fold increase in DSB numbers when ATM is missing. Note that the ratio of new hotspots to matched hotspots (shared between *Atm* null and *Atm* wt) is higher in the DSB-poor domain. Strengths of individual hotspots are not indicated (see main text). **(B)** Representative SPOll-oligo maps of a DSB-rich (left) and DSB-poor (right) domain at the distal end of chromosome 18 (data from ref.^17^). The subtelomeric region of chromosome 18 was reported to display frequent meiotic crossing over.^141^ In the absence of ATM, the sum of normalized SPOll-oligo counts increases more in the DSB-poor domain than in the DSB-rich domain, 2.4-fold and 1.3-fold respectively. This is because the existing weak hotspots in the DSB-poor domain increase more than the existing stronger hotspots in the DSB-rich domain, on average 5.8-fold in poor and 1.5-fold in rich domain. In addition, more new weak hotspots appear in the DSB-poor domain, such that the ratio of new to matched hotspots in the poor domain is higher (DSB-poor domain: 22 new to 10 matched hotspots; DSB-rich domain: 38 new to 34 matched hotspots). SPOll-oligo maps were smoothed with a 10001-bp Hann filter. RPM denotes reads per million. **(C)** Behavior of 5-Mb domains of chromosome 18 in relation to SPOll-oligo counts and DSB hotspot numbers in *Atm* wt and *Atm* null. (top left) DSB-poor domains (lower SPO11-oligo counts, colder domains) contain fewer hotspots. (top right) In the absence of ATM, strengths of domains with fewer hotspots in wild type tend to increase more. (bottom left) In the absence of ATM, normally colder domains tend to show more new hotspots relative to the number of matched hotspots in the domain. (bottom right) Domains with higher ratio of new to matched hotspots tend to be more strongly suppressed in the wild type by ATM. r represents Pearson’s correlation.

In budding yeast, Tel1 provides a more subtle contribution to genome-wide DSB patterning.^23^ Nevertheless, Tel1 absence causes changes in DSB distributions, as distinct subchromosomal domains (e.g., hotspots in subtelomeric or pericentric regions) accumulate fewer DSBs than the genome average early in meiosis. These alterations are not permanent, as later in meiosis DSB frequencies in the regions less sensitive to Tel1 loss level off at the genome average. Remarkably, this homeostatic compensatory behavior is implemented despite the increase in DSB numbers in *tel1*, and so it must be mediated by other regulatory systems, with this behavior showing the ability of meiotic cells to adjust to variations in DSB numbers.

Whole chromosomes provide an even larger size scale to evaluate DSB distribution. Smaller chromosomes typically experience more DSBs, which is attributed to a negative feedback mechanism tied to some feature of the engagement of homologous chromosomes.^56^ This evolutionarily conserved pathway negatively regulates DSB numbers by inhibiting further DSB formation on successfully synapsed chromosome segments.^10,56,108,109^ Perhaps smaller chromosomes take longer on average to engage with one another, allowing them to continue to undergo DSB formation for longer periods. Thus, in budding yeast and mouse, this behavior can be observed as a negative correlation of SPO11-oligo density with chromosome size.^16,17,56^ In *S. cerevisiae*, this anti-correlation is lost in mutants missing components of the homolog engagement pathway (“ZMM” mutants defective in homologous synapsis and recombination^110^). Similarly to *tel1*, these mutants produce more DSBs and display changes in DSB distributions.^56^ However, recent work in *S. cerevisiae* and mouse has reinforced the idea that homolog engagement and ATM/Tel1 represent distinct pathways for negative control of DSB formation^10^: In *Atm^-/^* and *tel1* mutants, SPO11-oligo density remains negatively correlated with chromosome size despite the increase in DSBs, although Tel is required for timely establishment of this relationship. Thus, chromosome-scale DSB patterns are not observably affected by ATM/Tel1.

ATM has an even more pronounced role in suppressing DSB formation on the non-homologous parts of the X and Y chromosomes. In wild-type spermatocytes, SPO11-oligo density in these regions is much lower than on autosomes, but in *Atm^-/-^*, these regions experience a greater increase in SPO11-oligo density than genome average.^17^ Although the molecular details remain to be elucidated, it is plausible that this suppression is needed to retain low numbers of DSBs in non-homologous sex-chromosome segments, as they are not competent for synapsis and meiotic recombination.

### A role for ATR in DSB control?

The more profound DSB distribution phenotype in *Atm^-/^* mice than *tel1* yeast may reflect the quantitatively more severe DSB number phenotype of *Atm^-/^* or the acquisition of different functions for the two proteins during meiosis. Redundancy with other factors must also be considered, especially in budding yeast, where redundancy between Tel1 and the Mec1 kinase (related to mammalian ATR) has been observed in other meiotic contexts. Both proteins phosphorylate Rec114, a SPO11 partner required for DSB formation, and Sae2.^26,97^ Moreover, both Tel1 and Mec1 play roles in DSB resection and directing DSB repair towards the homolog by inhibiting intersister recombination.^53,104,105^ While Tel1 controls the repair of low abundance DSBs occurring early in meiosis, it appears to become redundant with Mec1 for the repair of high abundance DSBs later in meiosis.^105^

If redundancy between Tell and Mecl exists in the control of DSB distribution, timing may be a key factor. For example, this could explain why the observed alterations in DSB patterning in the *tel1* mutant change over time in meiosis.^23^ We emphasize, however, that even though Mec1 may substitute for Tel1 to some extent or at specific times, Tel1 plays a dominant role in this process. This is supported by the fact that in the presence of Tell, the Mecl pathway is dispensable for the local control of DSB distribution.^20^

In mice, ATR does not appear to play a major role in regulating DSB numbers. SPOll-oligo complexes formed in spermatocytes with conditionally-deleted *Atr* are similar in number to those in wild type,^111^ while they are slightly increased in spermatocytes carrying a hypomorphic *Atr* allele.^112^ As in yeast, epistasis studies could bring new perspectives on the relationships of these two kinases. Redundancy is suggested in another context in that ATR partially substitutes for ATM in response to DSBs to phosphorylate H2AX in meiosis.^27,30^ ATR has an essential function in executing DSB repair in mouse meiosis, in addition to its previously described role in meiotic silencing of unsynapsed chromatin.^113,114^

## Mechanism of ATM/Tel1-dependent regulation of DSB formation

How ATM/Tel1 negatively regulates DSB formation is an active area of investigation. It is envisioned that ATM/Tel1 is activated by DSBs to phosphorylate proteins that regulate DSB formation. Kinase activity is critical, as a kinase-dead *tel1* mutation increases levels of Spoll-oligo complexes similarly to *tel1* null mutation.^23^ Similar experiments have not been performed in mice, however, because kinase-dead *Atm* mutations lead to embryonic lethality.^115,116^

Relevant phosphotarget(s) are not yet known, although REC114 and HORMAD1 are good candidates. REC114 is a conserved component of the DSB-promoting complex (REC114-MEI4-IHO1 in mouse, Rec114-Mei4-Mer2 in *S. cerevisiae;* see also ref.^117^), whereas HORMAD1 and its yeast ortholog, Hop1, are structural elements of meiotic chromosome axes that are required for normal DSB levels.^4,26,53,109,118–122^ Yeast Rec114 and Hopl are known Tell and Mecl targets, and they are phosphorylated in a DSB-dependent manner.^26,53,123^ Also, mouse HORMAD1 contain putative ATM/ATR phosphorylation sites^109,124^ and DSB formation promotes HORMAD1 phosphorylation.^124^

In yeast, experiments testing whether Tell-mediated DSB control operates via Rec114 phosphorylation have given ambiguous results. A *rec114* mutant lacking potential Tell and Mecl target SQ/TQ sites *(rec114-8A)* shows evidence of precocious and elevated DSB formation, and a putative phosphomimetic mutant *(rec114-8D)* shows reduced DSB levels.^26^ However, neither *rec114-8A* alone or in combination with a phosphomutant *hop1* phenocopies the elevated Spoll-oligo complex formation caused by Tel1 deficiency.^23^ Moreover, formation of multiple nearby DSBs on the same chromatid is not observed with non-phosphorylatable *rec114* and *hop1* alleles (V. Garcia and M. Neale, personal communication), unlike what is observed with the absence of Tell.^20^ These findings can be reconciled if there are multiple Tel1 targets (including Rec114 and Hop1), any of which is sufficient to inhibit DSB formation when phosphorylated.^23^

In mouse, so far, only one of the several HORMAD1 phosphorylation sites, Ser375, has been investigated.^113,124^ Ser375-phosphorylated HORMAD1 appears early in meiosis and it is associated with unsynapsed chromosomes throughout meiotic prophase.

However, whether phosphorylation at this site depends on ATM has only been examined at late meiotic stages, not the crucial early meiotic stages when negative feedback circuits are expected to regulate DSB formation.

It has been proposed that DSB-promoting complexes assemble on chromosome axes to which the chromatin loop is tethered and a DSB is formed^92,93,125,126^ (**Fig. 4**). Molecular details of this model integrating DSB formation with the loop-axis structure of meiotic chromosomes derive principally from data in yeast, but it has long been envisioned as an evolutionarily conserved process.^127^ Recent studies in mouse support this view.^95,118^ If correct, the tethering of chromatin loops at DSB sites would provide a structural framework within which ATM could phosphorylate members of the DSB-promoting complex and/or HORMAD1 to negatively regulate DSB formation and thus locally shape DSB distribution and hotspot usage (**Fig. 4**).

**Figure 4.**
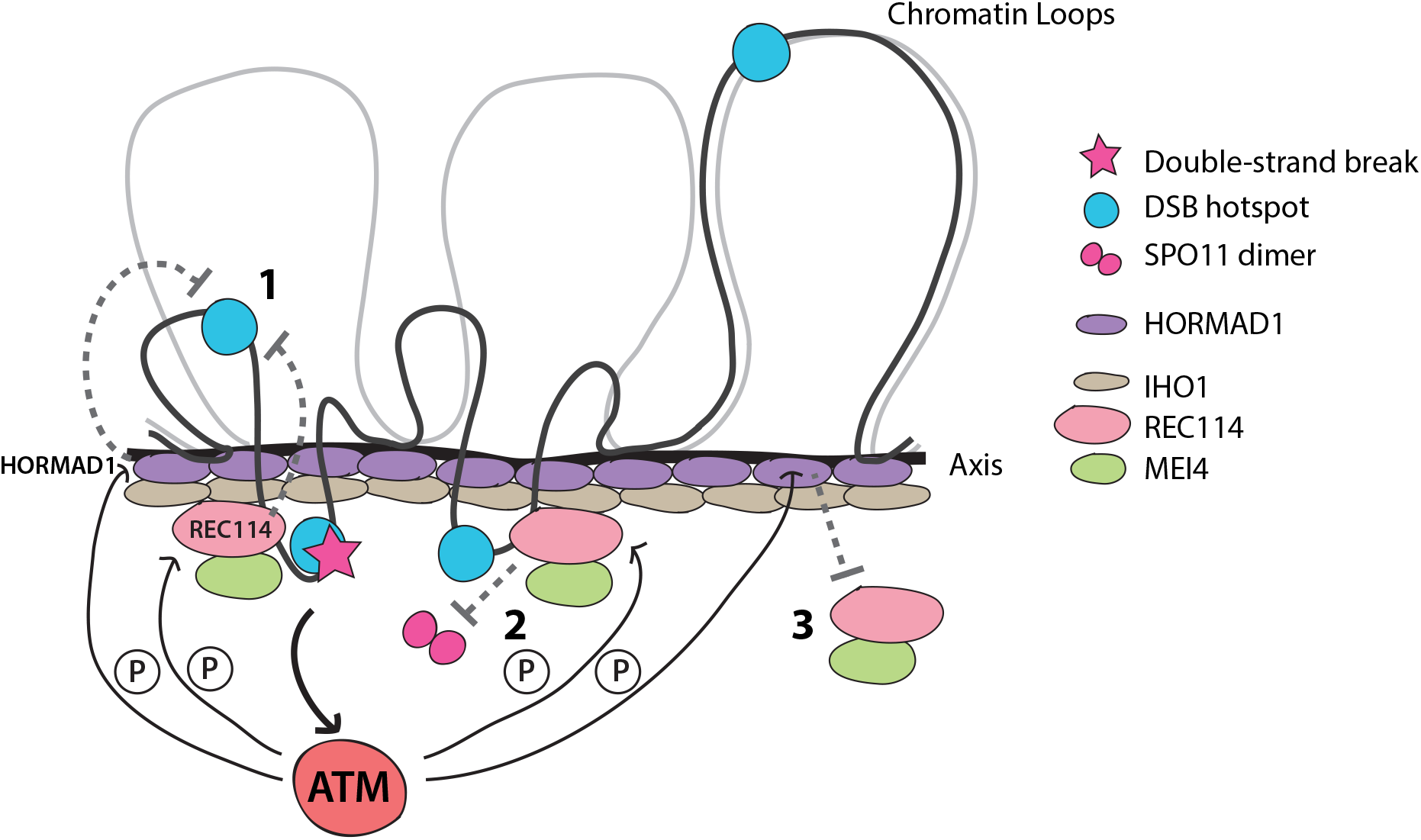
Hypothetical model for ATM-mediated DSB control. The IHO1-REC114-MEI4 complex (DSB-promoting complex) and the axis-structural component HORMAD1 promote DSB formation by SPO11. HORMAD1 interacts with IHO1. A DSB is formed preferentially at a DSB hotspot, in a chromatin loop tethered to the axis. In response to the DSB, ATM is activated and phosphorylates REC114 and/or HORMAD1. Phosphorylation events separately or in combination inhibit additional DSB formation at the same chromatin loop (1) or at the adjacent loop pre-activated for DSB formation (2), by inhibiting or destabilizing of the IHO1-REC114-MEI4 complex. (3) HORMAD1 phosphorylation in the vicinity of the DSB prevents the assembly of the IHO1-REC114-MEI4 complexes and thus DSB formation.

If tethering of chromatin loops explains local effects of ATM, how might ATM influence DSB distributions at larger size scales? On the assumption that the relevant ATM targets are the same for both local and large-scale DSB control, it has been proposed that the differential susceptibility of Mb-sized chromosome domains to ATM-mediated DSB suppression reflects variability in the amount of ATM substrates (e.g., REC114 and/or HORMAD1) along the chromosome axis.^17^ In this view, the underlying density of these factors may determine domains of differing intrinsic DSB-forming potential as well as of differential sensitivity to ATM. In budding yeast, DSB-rich genomic regions are enriched for Hop1 and Rec114.^126^ These domains of enrichment also tend to be more responsive to the Tel1-independent negative feedback regulation of DSBs attributed to homolog engagement.^56^

In mice, there is an inverse correlation for Mb-sized domains between the SPO11-oligo density in wild type and the fold-increase in SPO11-oligo density in the absence of ATM.^17^ That is, relatively DSB-poor domains in wild type also tend to be the domains that are more highly suppressed by ATM (also see Fig. 3). Thus, studies in mice of correlations between the density of potential ATM substrates and the strength of ATM-dependent regulation are likely to be informative. Intuitively, domains enriched for substrates might be more responsive to ATM (positive correlation), but the opposite can be also imagined (negative correlation). In particular, domains occupied by fewer substrates might be more responsive if the ATM proficiency is measured by how “good” ATM is in phosphorylating all of its available targets. In this case, fewer targets would simply mean less “work” for ATM.

A mentioned above, histone H2AX is also a target for ATM-mediated phosphorylation at residue 139 (γH2AX).^128^ Because γH2AX is not spatially restricted to DSB sites but spreads from DSBs to surrounding chromatin^129^, ATM-dependent γH2AX spreading could theoretically function to suppress DSB formation in the vicinity of the initial break. The extent of γH2AX-positive domains in meiosis has been not assessed, but in mitotically cycling cells, γH2AX extends for up to 1-2 Mb in mammals^130^ and for up to 50 kb in *S. cerevisiae.^131^* However, although this idea is appealing, *H2ax*-null mice do not phenocopy *Atm^-/-^*.^132,133^ Also, in *S. cerevisiae, h2a* mutants do not show DSB clustering like *tel1* (V. Garcia and M. Neale, personal communication, and ref.^20^). Thus, H2AX is not critical for ATM-regulated negative feedback. Possible redundancy should also be considered in this case, however, since at least one other Tell/Mecl-dependent histone phosphorylation (Thr 129 phosphorylation on H2B) has been identified in vegetative yeast.^134^

## Consequences of the lack of ATM

While the grossly elevated number of DSBs in Atm-null meiosis is presumably a major source of meiotic defects,^22^ alteration in DSB distribution is likely to be as critical.^17^ As discussed above, loss of local DSB suppression in *Atm^-/-^* may provoke SPO11 to cut multiple chromatids at the same time, which would damage the repair template, or to cut multiple times at adjacent genomic positions, which may be problematic to repair. It is interesting to consider how loss of control of local DSB patterning contributes to the overall increase in DSB numbers. *Atm^-/-^* spermatocytes have a relatively small increase in cytological foci of DMC1 (the meiosis-specific paralog of the RAD51 strand-exchange protein that assembles on resected strands) compared with the increase in SPOll-oligo complexes;(ref.^22^ and A.L., J.L., S.K., M.J., unpublished observations) and similar observations have been reported previously for RAD51 foci in *Spo11*^+/-^*Atm^-/-^* spermatocytes^51^ suggesting that fewer foci than expected may mark multiple, closely spaced DSBs. An alternative is that fewer DMC1/RAD51 arise because of the involvement of ATM in other functions, such as resection, as seen in mitotic cells^135–137^ and in yeast meiosis,^104,105^ and/or interhomolog bias,^53^ which may cause more rapid intersister DSB repair. If ATM has these functions in meiosis, like Tel1 in budding yeast, their loss might exacerbate the consequences of dysregulated DSB formation.

Other consequences of the lack of ATM should also be considered that were not previously anticipated. For example, extra DSBs formed in DSB-cold domains may present issues for repair if the chromatin composition or chromosome structure of cold domains limits access of the DSB repair machinery. Although rather speculative at this point, chromatin context is known to affect DSB repair pathway choice in mitotic cells.^138^ Enrichment for meiosis-specific factors in different domains could also affect recombination pathway choice, for example, as proposed for differential Zip3 or Hop1 occupancy in yeast.^139,140^ It will be interesting to determine whether changes in recombination outcomes observed in the absence of ATM/Tell^24,51^ are a direct consequence of large-scale changes in DSB distributions.

## Concluding remarks

The number of DSBs and their distribution in meiosis is highly regulated and the ATM kinase critically contributes to this complex process. While clearly important for a successful meiosis, ATM control of DSB formation also has the power to shape genome evolution. Understanding the underlying mechanisms is a key future priority. Deciphering relevant ATM targets and mapping their positions in the genome relative to SPOll-oligo maps will shed light onto how ATM molds the DSB landscape. Moreover, correlating SPO11 activity across the genome with other cellular processes (e.g., transcription), chromatin modifications, and meiotic chromosome architecture (loop-axis organization) may reveal interesting relationships. Conversely, it could be fruitful to explore changes in chromatin dynamics in the absence of ATM. A detailed investigation of DSB repair in the absence of ATM at distinct DSB sites and genome-wide will be crucial for understanding the effect of ATM-mediated DSB control in the suppression of aberrant meiotic events.

## Acknowledgments

We thank members of the Jasin and Keeney labs for helpful discussions.

## Disclosure of potential conflicts of interest

No potential conflicts of interest were disclosed.

## Funding

MSKCC core facilities are supported by NIH grant P30 CA008748. This work was supported by March of Dimes grant 1-FY17-799 (M.J.), NIH grant R35 GM118092 (S.K.), and NIH grant R35 GM118175 (M.J.).

